# An EpCAM-Targeted Mirror-Image DNA Nanostructure for Precise Drug Delivery in Triple-Negative Breast Cancer

**DOI:** 10.64898/2026.06.11.730265

**Authors:** Sihan Wu, Terry Farkaly, Wen Zhang

## Abstract

Triple-negative breast cancer (TNBC) remains a major therapeutic challenge due to the lack of effective molecular targets and the dose-limiting off-target toxicity of conventional chemotherapy. Here, we design and construct a mirror-image DNA (L-DNA) nanostructure functionalized with an epithelial cell adhesion molecule (EpCAM)-specific aptamer for targeted delivery of doxorubicin (DOX) to TNBC cells. The L-DNA nanostructure retains thermodynamic properties comparable to natural D-DNA while exhibiting substantially enhanced resistance to nuclease and serum-mediated degradation due to its mirror-image chirality. Thermal melting and serum stability assays confirmed superior structural stability of the L-DNA nanostructure compared to D-DNA counterparts. In vitro cytotoxicity studies demonstrated that the EpCAM-targeted L-DNA nanostructure has the potential to selectively inhibit the growth of EpCAM-positive TNBC cells while reducing cytotoxicity in normal cells. These findings demonstrate that combining aptamer targeting with mirror-image DNA nanotechnology provides a stable and selective nanoplatform for chemotherapeutic delivery, which can potentially improve the precision and therapeutic efficacy of treatment for aggressive breast cancers.

## INTRODUCTION

Triple-negative breast cancer (TNBC) is a highly aggressive subtype of breast cancer defined by the absence of estrogen receptor (ER), progesterone receptor (PR), and HER2 expression. TNBC accounts for approximately 10 - 20% of all breast cancer cases and is associated with the poorest prognosis compared with hormone receptor–positive and HER2-positive subtypes^1^. Globally, breast cancer causes nearly half a million deaths each year, of which an estimated 150,000 are attributable to TNBC, reflecting the lack of effective targeted therapies for this disease^1^. Doxorubicin (DOX) remains one of the most commonly used chemotherapeutic agents for TNBC. However, its clinical application is limited by poor tumor selectivity, systemic toxicity, and cumulative cardiotoxicity^2^. Consequently, developing more precise drug-delivery systems has become an important goal in cancer treatment.

Conventional chemotherapy is frequently limited by insufficient tumor selectivity and severe systemic toxicity. To address these challenges, a variety of nanoscale delivery platforms have been explored to improve drug accumulation in tumors while minimizing damage to healthy tissues^3^. Existing systems such as lipid nanoparticles, viral carriers, and synthetic polymers have shown promise in cancer therapy; however, their broader application is often constrained by issues including immunogenic responses, formulation complexity, and limited cargo capacity. Among emerging nanomaterials, nucleic acid-based nanostructures have gained increasing interest because they can be precisely programmed through sequence-directed self-assembly^4-6^. The predictable base-pairing properties of nucleic acids enable the construction of structurally defined architectures with tunable sizes, shapes, and functionalities. In particular, nucleic acid nanostructures have been widely investigated as delivery vehicles due to their biocompatibility, structural flexibility, and ability to incorporate therapeutic modules into a single scaffold^7^. These nanoscale assemblies can also facilitate cellular uptake and molecular recognition, making them attractive candidates for precision drug delivery.

Despite their advantages, the clinical translation of nucleic acid nanotechnology remains challenging because native nucleic acid molecules are intrinsically unstable under physiological conditions^8^. Nucleases are highly abundant in serum and tissues, leading to rapid degradation of nucleic acid nanoparticles and short in vivo persistence^9, 10^. Therefore, alternative nucleic acid systems with inherently improved biostability are needed for the development of more durable therapeutic nanostructures.

Mirror-image nucleic acids represent one such strategy. L-nucleic acids are composed of enantiomeric nucleotides with opposite chirality to naturally occurring D-nucleic acids^11^ as depicted in Figure 1A. Because naturally evolved nucleases cannot efficiently recognize these mirror-image structures, L-RNA and L-DNA exhibit remarkable resistance to enzymatic degradation^12, 13^. Importantly, mirror-image nucleic acids retain the ability to form predictable secondary and tertiary structures and can be further functionalized through chemical modification^14-17^. Previous studies have demonstrated the utility of L-nucleic acids in a range of biomedical applications, including Spiegelmer-based targeting ligands^18-21^, therapeutic L-nucleosides^13^, and mirror-image DNA nanostructures for cargo delivery^22, 23^. More recently, structurally sophisticated L-DNA materials, such as pH-responsive i-motif assemblies^24^ and biostable DNA hydrogels^25^, have further illustrated the versatility of mirror-image nucleic acid engineering. These developments suggest that L-nucleic acid nanotechnology may provide a powerful foundation for constructing stable and functional delivery systems for therapeutic applications.

**Figure 1.**
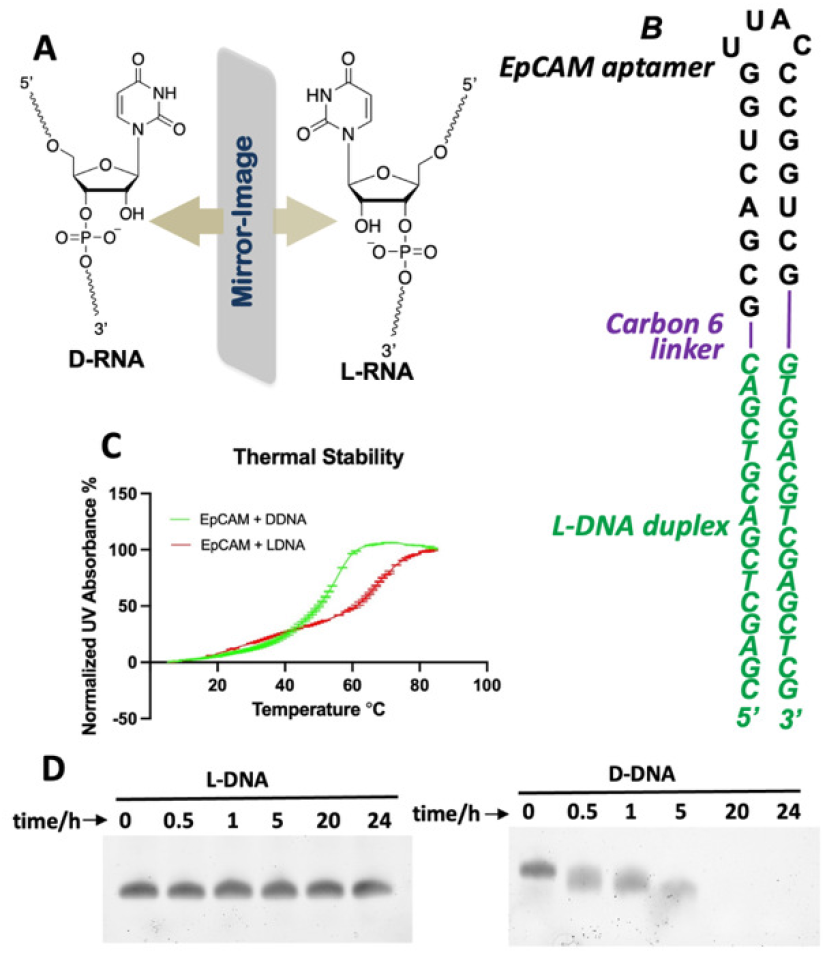
Design, construction, and characterization of L-DNA nanostructure. (A) Chemical structures of D- and L-ribonucleotides; (B) Sequence and secondary structure of synthetic L-DNA nanostructure. Green: L-DNA duplex for DOX loading; purple: carbon 18 linker; black: EpCAM aptamer. (C) *T*_m_ melting curves for the assembly of the aptamer-L-DNA structure (red) and aptamer-D-DNA structure (green). (D) Stability assay of L- (left) and D-DNA (right) after treating with phosphodiesterase.

On the other hand, the epithelial cell adhesion molecule EpCAM (CD326) is a transmembrane glycoprotein involved in epithelial cell adhesion and intracellular signalling^26^. Although EpCAM is expressed at low levels in normal epithelial tissues, it is overexpressed in many solid tumors, including a substantial proportion of TNBC cases^27^. EpCAM overexpression in TNBC has been associated with poor prognosis, lymph node metastasis, and decreased survival, making it an attractive therapeutic target^27^. Therefore, we decide to conjugate an EpCAM targeting aptamer to our L-DNA molecule for the enhanced specifical drug delivery. Aptamers are short single stranded DNA or RNA molecules that are selected through systematic evolution of ligands by exponential enrichment (SELEX) and have emerged as powerful targeting ligands due to their high specificity, small molecular size, and low immunogenicity^28, 29^. Compared with monoclonal antibodies, aptamers provide improved tissue penetration and lower production variability. Aptamers targeting EpCAM have been developed and shown to bind selectively to EpCAM-positive cancer cells, enabling targeted therapeutic delivery^30^.

## MATERIALS AND METHODS

### Preparation of EpCAM Aptamer, EpCAM-D-DNA and EpCAM-L-DNA Oligonucleotides

The L-DNA containing oligonucleotides used for biochemical and cellular studies were solid-phase synthesized using ABI 394 DNA/RNA synthesizer. The phosphoramidite compounds were obtained from Chemgenes or Glen Research, Inc. The DMTr-on in-house synthesized oligonucleotides were deprotected by AMA (1:1 mixture of ammonium hydroxide and methyl amine solution) for 15 min at 65°C, followed by the Et_3_N•3HF treatment; and purified by denaturing PAGE gel electrophoresis. After purification, the oligonucleotides were desalted and re-concentrated to the proper concentrations for nanostructure construction and biochemical experiments.

All the sequence information and the MS characterization are listed in Supporting Information, with the designed structure of the EpCAM-targeted L-DNA nanostructure illustrated in Figure 1B. All the nanostructures were self-assembled by mixing the different strands at the same molar concentrations in TMS buffer (25 mM Tris-HCl pH 7.5, 10 mM MgCl_2_, 50 mM NaCl) or PBS buffer (137 mM NaCl, 2.7 mM KCl, 10 mM Na_2_HPO_4_, 2 mM KH_2_PO_4_, pH 7.4), heated to 90°C for 2 min and slowly cooled to 23 °C.

### L-DNA enzymatic stability assays

Both L- and D-nucleic acids were incubated in 25% phosphodiesterase II at 37°C for different times. 200 ng of DNA were taken at 0 min, 30 min, 1 hr, 5 hr, 20 hr and 24 hr time points, immediately mixed with 6× native loading buffer and analyzed by 10% native PAGE in TB buffer. The gels were stained with Sybr Gold (Invitrogen) and washed briefly before imaging using the Bio-rad ChemiDoc Imaging System.

### Melting temperature measurement of L-DNA nanostructures

Melting temperature measurement was performed using the Varian Cary 300 UV-Visible Spectrophotometer equipped with Agilent temperature controller. The experiments were performed using the samples (0.2 µM RNA or DNA structures) dissolved in water. The samples were heated to 85ºC and allowed to cool down to room temperature slowly. The UV absorbance at 260 nm was recorded at a heating rate of 0.5 ºC/min from 5 °C to 85 °C.

### Cell culture

MDA-MB-231 (RRID: CVCL_0419) and HEK-293T (RRID: CVCL_0063) cells were purchased from the American Tissue Type Collection (ATCC, Manassas, VA), or gift from Indiana University Melvin and Bren Simon Comprehensive Cancer Center. Cells were maintained in DMEM (Corning, Cat. #10-013-CV) supplemented with 10% FBS. All cell lines were maintained at 37°C in a fully humidified atmosphere containing 5% CO_2_, used at early passage for experiments, and tested to be free of mycoplasma contamination.

### Fluorescent microscope analysis

MDA-MB-231 cells were seeded onto poly-D-lysine–coated coverslips in 24-well plates at a density of 1 × 10^5 cells per well and cultured overnight in DMEM supplemented with 10% fetal bovine serum at 37 °C in a humidified atmosphere containing 5% CO_2_. Cells were incubated with Cy5-labeled EpCAM-targeted L-DNA nanostructures or Cy5-labeled scramble oligo lacking the EpCAM aptamer at a final concentration of 50 nM for 2 h in complete culture medium. Following incubation, cells were washed three times with phosphate-buffered saline (PBS) to remove unbound nanostructures and fixed with 4% paraformaldehyde for 15 min at room temperature. Cell nuclei were stained with DAPI for 5 min prior to imaging.

Fluorescence images were acquired using an inverted fluorescence microscope equipped with DAPI and Cy5 filter sets. Images were processed using ImageJ software under identical exposure conditions for all experimental groups.

### Flow cytometry analysis

Flow cytometry was performed to quantify the internalization of Cy5^+^, aptamer^+^ and Cy5^+^, aptamer^−^ nanostructures. MDA-MB-231 cells were cultured in DMEM with 10% FBS at 37°C in a 5% CO_2_ incubator until they reached 80% confluence. Cells were seeded in 6-well plates at a density of 1 × 10^5^ cells per well and allowed to adhere overnight. Nanostructures were added at a final concentration of 50 nM in opti-MEM medium and incubated with cells for 4 hours. After incubation, cells were washed three times with PBS, detached and resuspended in 1 mL PBS for analysis. Flow cytometry was performed with Cy5 fluorescence detected and at least 10,000 events were collected for each sample. Data were analyzed using FlowJo software (FlowJo LLC), and the data are presented as the percentage of the cells.

### Formation of DOX/nanostructure complexes

Doxorubicin was loaded into the EpCAM–L-DNA nanostructure via spontaneous binding with double-stranded L-DNA domains. Briefly, pre-assembled EpCAM–L-DNA nanostructures (2 μM final concentration) were incubated with DOX at varying molar ratios (DOX:L-DNA base pair ratios of 1:1) in loading buffer: 50 mM NaCl, 10 mM phosphate buffer, pH 7.4, 10 mM MgCl_2_). The mixture was incubated at room temperature for 2 h under gentle agitation to allow intercalation equilibrium. Unbound DOX was removed by centrifugal ultrafiltration using 10 kDa molecular weight cutoff filters (Amicon Ultra, Millipore) and washed three times with a loading buffer. The retained nanostructure was collected and stored at 4 °C until further use. For comparison, DOX loading into analogous EpCAM–D-DNA nanostructures was performed under identical conditions.

### In Vitro Cytotoxicity Assessment by Alamar Blue Assay

Cell viability was evaluated using the Alamar Blue assay^31^ (Invitrogen, Cat. No. A50101) according to the manufacturer’s protocol. MDA-MB-231 and HEK-293T cells were seeded in poly-D-lysine–coated 96-well plates at a density of 8,000–10,000 cells per well in 100 μL of complete growth medium and allowed to adhere overnight at 37 °C in a humidified atmosphere containing 5% CO_2_. Cells were treated with serial dilutions of the following formulations: free doxorubicin (DOX), aptamer–L-DNA–DOX nanostructures, aptamer–D-DNA–DOX nanostructures, and phosphate-buffered saline (PBS) as a negative control. Drug-containing solutions were prepared in complete medium and added to cells to achieve a final treatment volume of 100 μL per well. Treatments were administered for 3 h, after which the medium was removed and replaced with fresh complete medium. Subsequently, after, 10 μL of alamarBlue reagent (10% v/v final concentration) was added to each well. Cells were incubated for 1 h at 37 °C, and fluorescence was measured using a Synergy H1 microplate reader (BioTek Instruments) with excitation at 560 nm and emission at 590 nm. Cell viability was calculated as the percentage of fluorescence intensity relative to untreated control cells.

## RESULTS

### Stability of L-DNA nanostructure

The thermal stability of the nanostructures were determined by measuring the UV absorbance at 260 nm as a function of temperature. The normalized result is shown in Figure 1C, which indicates the annealing transition of different complexes of oligo assemblies. Both aptamer-D-DNA and aptamer-L-DNA complexes exhibited characteristic melting profiles corresponding to the transition from double-stranded to single-stranded structures. As temperature increased, a gradual rise in UV absorbance was observed due to the hyperchromic effect associated with nucleic acid strand separation. The aptamer-D-DNA nanostructure displayed a melting transition at approximately 58–62 °C, while the aptamer-LDNA nanostructure showed a slightly higher melting temperature of approximately 70–75 °C. The rightward shift of the melting curve for aptamer-L-DNA indicates improved thermal stability compared with the conventional D-DNA counterpart.

These results demonstrate that the mirror-image L-DNA nanostructure maintains strong structural integrity over a broader temperature range. The enhanced stability is consistent with the physicochemical properties of mirror-image nucleic acids, which preserve Watson–Crick base pairing while exhibiting altered chirality. This increased thermal stability further supports the use of L-DNA nanostructures as robust scaffolds for nucleic acid–based drug delivery systems under physiological conditions.

It has been known about the enzymatic stability of L-nucleic acid under physiological conditions^32^. We then attempted to evaluate the enzymatic stability of our designed D-DNA and L-DNA nanostructures for drug delivery. By incubating the samples in 25% phosphodiesterase II conditions and analyzing degradation over time, as shown in Figure 1D, the L-DNA nanostructure remained largely intact over the entire 24 h incubation period, with minimal reduction in band intensity and no detectable fragmentation. In contrast, the D-DNA nanostructure exhibited rapid degradation, with a significant decrease in the intact band observed within the first 1–5 h, accompanied by the appearance of smeared and lower-molecular-weight fragments indicative of nuclease-mediated cleavage. By 20 h, the D-DNA nanostructure was almost completely degraded under the same conditions.

These results demonstrate that the L-DNA nanostructure possesses significantly enhanced resistance to nuclease-mediated degradation compared with D-DNA. The improved stability is likely due to the mirror-image chirality of L-DNA, which prevents recognition and cleavage of the whole nucleic acid construct by natural nucleases. This enhanced enzymatic stability supports the potential of L-DNA nanostructures as robust platforms for nucleic acid–based drug delivery systems.

### Specific cancer cell targeting by EpCAM aptamer-conjugated L-DNA nanostructure

To facilitate the successful administration of anti-cancer agents into cancer cells, our designed nanomedicine must first bind to the receptor on cancer cell membrane, and penetrate the cell membrane, allowing for the subsequent release of therapeutics into the cytoplasm. To validate the efficient binding and targeting of the nanostructures to cancer cells, fluorescent microscopy and flow cytometry experiments were performed. Media containing 50 nM aptamer^+^, Cy5^+^ nanostructure was used to treat the TNBC cell line MDA-MB-231. For comparison, the same incubations were performed using the Cy5 labelled scrambled oligo without the aptamer (Cy5^+^, aptamer^-^ structure), to validate the specific targeting. Following the nanostructure incubation for 2h and washing steps, the cells were fixed, the nuclei were stained with DAPI.

Microscopy images show increased fluorescence from Cy5^+^, aptamer^+^ nanostructure following incubation with MDA-MB-231 cells (Figure 2A). The observed Cy5 signals were surrounding the stained nuclei, suggesting the nanostructures were internalized and delivered to cytoplasm of the cancer cells. In comparison, the Cy5^+^, aptamer^-^ oligo was not binding or delivered into cancer cells, due to the lack of aptamer to direct cell targeting. This successful detection of our designed nanostructure within cancer cells holds great promise for the potential to administer therapeutics with a high degree of stability and precision.

**Figure 2.**
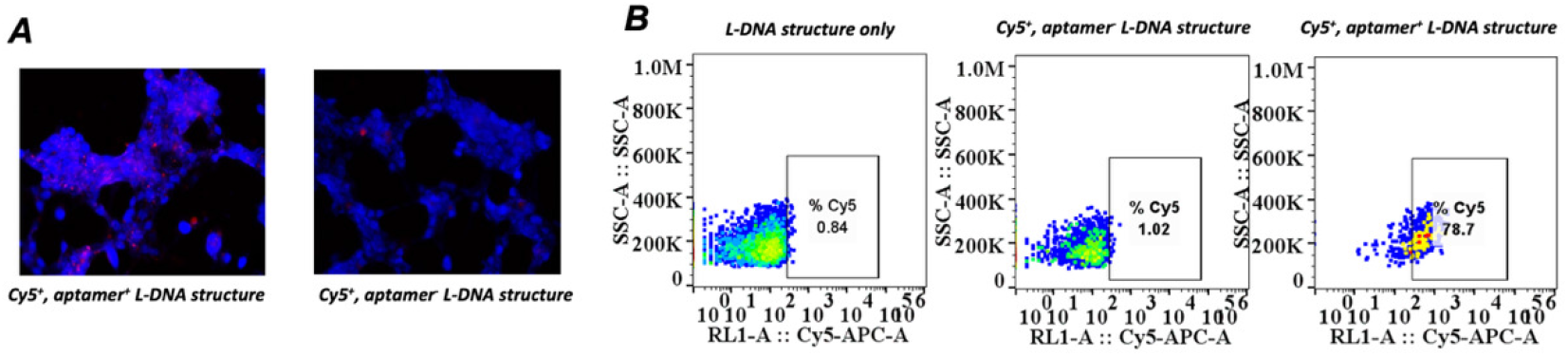
Binding and targeting of L-DNA nanostructure mediated by the EpCAM aptamer. (A) Fluorescent microscope images indicating the binding of the nanostructure to MDA-MB-231 cells (blue: nuclei; red: L-DNA nanostructures, the concentration of each fluorescent RNA nanostructures in treatment is 50 nM). (B) Flow cytometry analysis of Cy5-positive cells after treatment with Cy5^+^, aptamer^+^ L-DNA and Cy5^+^, aptamer^−^ L-DNA structures. The presence of EpCAM aptamer significantly increased the efficiency of nanostructure’s binding to cancer cells, with 78.7% of cells Cy5-positive in the aptamer^+^ condition compared to 1.02% in the aptamer^−^ condition.

Moreover, flow cytometry was conducted to quantify the internalization efficiency of Cy5^+^, aptamer^+^ and Cy5^+^, aptamer^−^ nanoparticles to the cancer cells. The results showed that 78.7% of cells were Cy5-positive when treated with aptamer^+^ nanoparticles, compared to only 1.02% for aptamer^−^ nanoparticles (Figure 2B). This dramatic difference demonstrates that the presence of EpCAM aptamer significantly enhances receptor-mediated uptake of the nanostructure. These findings confirm the selective targeting of aptamer^+^ nanostructure via EpCAM-mediated endocytosis, consistent with the fluorescence microscopy observations.

### Delivery of L-DNA nanostructures to cancer cells for viability assay

The cytotoxic effects of free DOX, aptamer^+^ DOX/D-DNA conjugate, and aptamer^+^ DOX/L-DNA conjugate were evaluated in MDA-MB-231 breast cancer cells and HEK293T normal cells using a cell viability assay. Cells were treated with increasing concentrations of DOX (4-1000 nM for cancer cells and 2-250 nM for normal cells), and cell viability was measured relative to the control (Figure 3).

**Figure 3.**
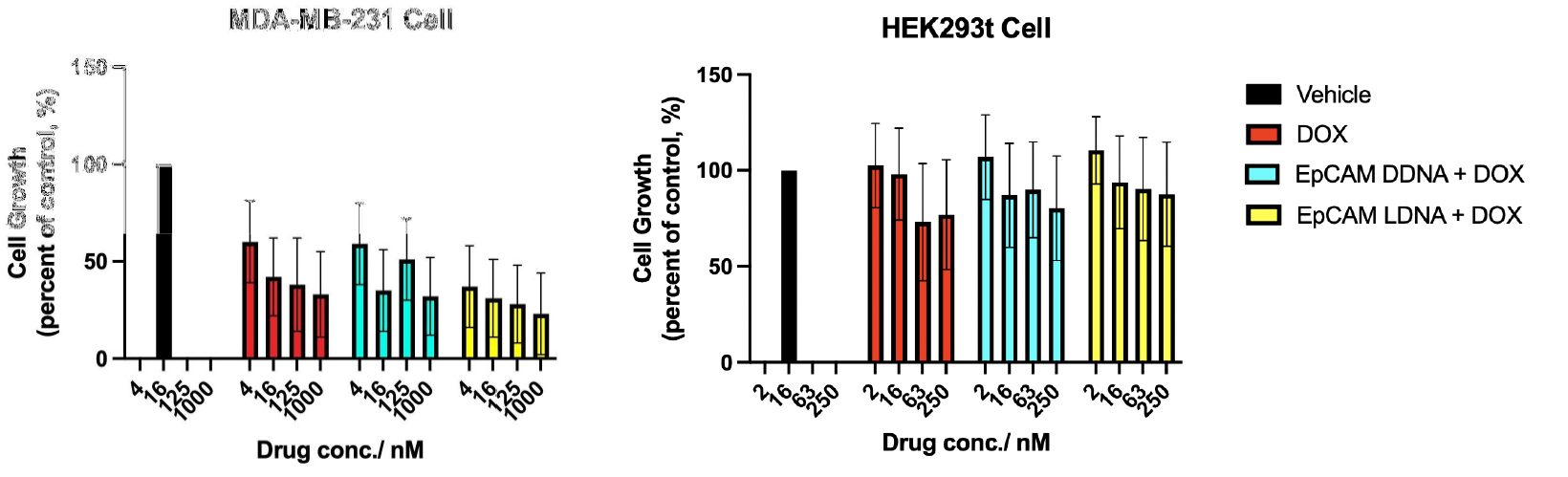
Comparison of inhibition of cell growth in MDA-MB-231 and HEK293t cells by nanostructures. (A) Alamar blue assay using different nanostructures to treat MDA-MB-231 at 4 different concentrations. (B) Alamar blue assay using different nanostructures to treat HEK293t cells at 4 different concentrations. The cell growth inhibitory effects were compared between different nanostructures and different concentrations. Data are representative of triplicate experiments. Error bars represent the mean ± SEM from three independent experiments (n = 3).

In MDA-MB-231 cells, all DOX-containing treatments reduced cell viability in a dose-dependent manner (Figure 3A). Both D-DNA+DOX and L-DNA+DOX nanostructures exhibited comparable or slightly enhanced cytotoxic effects compared with free DOX. At higher drug concentrations, L-DNA+DOX showed the strongest inhibition of cancer cell growth, suggesting efficient delivery of the chemotherapeutic payload to EpCAM-positive tumor cells. In contrast, in HEK293T normal cells, the nanostructure-mediated treatments exhibited slightly reduced cytotoxicity compared with free DOX, especially at 63 and 250 nM concentrations. This maintenance of high viability possibly indicates decreased off-target toxicity. This selective effect suggests that EpCAM-mediated targeting improves therapeutic specificity by preferentially affecting cancer cells while sparing normal cells.

These results demonstrate that the EpCAM-functionalized nucleic acid nanostructures enable effective delivery of DOX to EpCAM-expressing cancer cells while reducing cytotoxic effects on normal cells. Consistent with previous findings reported for mirror-image nucleic acid nanostructures, the L-DNA platform provides a stable and efficient delivery vehicle for targeted cancer therapy.

## DISCUSSION

The present study demonstrates that mirror-image DNA nanostructures can serve as stable and functional platforms for targeted chemotherapeutic delivery in triple-negative breast cancer. By integrating an EpCAM-targeting aptamer with an L-DNA scaffold, we generated a nanostructure capable of selectively delivering doxorubicin to cancer cells while exhibiting substantially improved resistance to thermal and enzymatic degradation compared with conventional D-DNA systems. These findings highlight the unique advantages of mirror-image nucleic acids for biomedical applications in which structural stability under physiological conditions is essential. As the major challenge in nucleic acid nanotechnology is the rapid degradation of natural nucleic acids in biological environments, our results further support the concept that chirality engineering provides an effective strategy to overcome this limitation. The observed reduction in cytotoxicity toward noncancerous cells also suggests that combining molecular targeting with highly stable nanostructures may improve the therapeutic selectivity of conventional chemotherapeutic agents.

It is important to note that the effectiveness of EpCAM-targeted delivery also depends on the level of EpCAM expression across different tumors. Since TNBC exhibits substantial molecular heterogeneity, future translational applications of this strategy would likely require biomarker-based patient stratification to identify tumors with sufficient EpCAM expression for effective targeting. In addition, the current work was limited to in vitro characterization, and further in vivo studies will be necessary to evaluate pharmacokinetics, biodistribution, tumor retention, systemic toxicity, and therapeutic efficacy in animal models.

## CONCLUSION

In summary, we developed an EpCAM-targeted L-DNA nanostructure for the targeted delivery of doxorubicin to TNBC cells. The mirror-image DNA scaffold demonstrated strong thermal stability and significantly improved resistance to enzymatic degradation compared with conventional D-DNA nanostructures. The microscopic imaging and flow cytometry results suggest the specific targeting of the constructed nanostructure to cancer cells. In vitro cytotoxicity experiments further showed that the EpCAM-targeted nanostructure effectively inhibited the growth of EpCAM positive cancer cells while reducing toxicity toward normal cells. Overall, this work demonstrates that mirror-image DNA nanotechnology represents a promising platform for the development of stable and selective nucleic acid based drug delivery systems.

## SUPPORTING INFORMATION

Additional experimental details, materials and methods, including sequences of the synthetic DNAs and their MS characterizations.

## ACKNOWLEDGEMENTS

I thank Zhang lab members for helpful discussions, insightful commentary, and careful revision of the manuscript. We thank Dr. J. Meng and the Chemical Genomics Core at IUSM for spectropolarimeter analysis.

## TOC Graphic

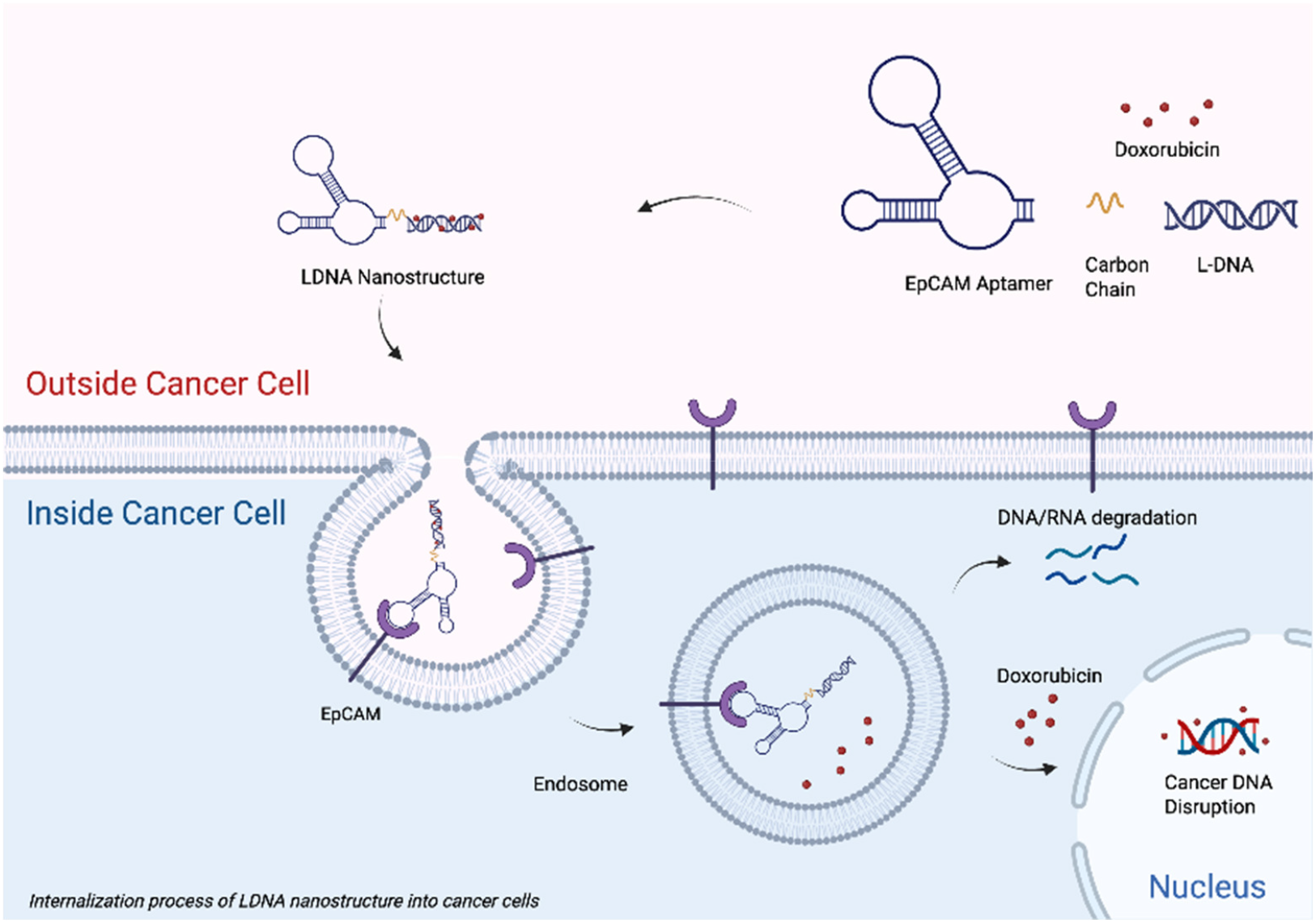

