## Supporting Information for "An EpCAM-Targeted Mirror-Image DNA Nanostructure for Precise Drug Delivery in Triple-Negative Breast Cancer"

**CONTENTS**

1. General methods S2

2. List of sequences of all the oligonucleotides and MS characterizations S3

#### 1. General Methods.

**1a. General considerations.** All chemicals were purchased from Fisher Scientific, VWR or Sigma-Aldrich unless otherwise noted. All the nucleoside phosphoramidites for solid-phase synthesis were from Chemgenes Corporation or Glen Research. All oligonucleotides, including fluorophore modified RNAs, 2'-F-modified RNAs, alkyne-modified RNAs and RNA-DNA chimeric strands, were synthesized in house. The alkyne modified phosphoramidintes were purchased from Hongene Biotech.

**1b. RNA oligonucleotides synthesis.** RNA oligonucleotides were synthesized by standard solid-phase phosphoramidite chemistry on a ABI RNA/DNA oligonucleotide synthesizer. Cleavage and elution of the full-length products from 1 µmol universal CPG-solid support columns, as well as removal of protecting groups on the nucleobases and phosphates, was carried out by heating the solid support for 15 min at 65 °C. The resultant homogeneous mixtures were first concentrated under reduced pressure for 2 hrs on a Genevac EZ-2 table top speedvac system (Genevac, Stone Ridge, NY), then lyophilized to dryness on a Labconco Benchtop 4.5 L freeze-drier (Labconco, Kansas City, MO) at <200 mTorr overnight to afford off-white solid residues. The residues were then resuspended in 115 µL of DMSO. 75 µL of TEA and 65 µL of TEA·3HF were added, and the solutions were heated for 2.5 hr at 65 °C to remove the TBDMS protecting group on the ribose 2'-hydroxyl group. The mixtures were homogeneous and pale to golden yellow in color. After cooling to room temperature (~30 mins), the RNA sample was desalted and detritylated by using the Glen-Pak RNA purification cartridge, following the protocol from Glen Research Inc. The purification of the desired products was performed by running denaturing polyacrylamide gels (urea-PAGE). The gel pieces containing the RNA fractions were collected, fragmented, and soaked in water for 12 hours. The RNA samples were then desalted by RNA precipitation using sodium chloride and ethanol. The precipitates were spun down (12000 rpm, 40 mins) and supernatants were removed by decanting. The resulting white solids were washed twice with absolute ethanol; the samples were then dried under high vacuum overnight before dissolving into water for appropriate concentrations.

#### 2. List of sequences of all the oligonucleotides

| entry | Sequence (5′ →3′) |
|---|---|
| EpCAM aptamer | GCGACUGGUUACCCGGUCG |
| EpCAM-L-DNA | **CGAGCTCGACGT**-Carbon6-<br>GCGACUGGUUACCCGGUCG-Carbon6<br>**ACGTCGAGCTCG** |
| Cy5 labeled<br>EpCAM-L-DNA | Cy5-**CGAGCTCGACGT**-Carbon6-<br>GCGACUGGUUACCCGGUCG-Carbon6<br>**ACGTCGAGCTCG** |
| EpCAM-D-DNA | CGAGCTCGACGTGCGACUGGUUACCCGGUCG<br>ACGTCGAGCTCG |

### underlined nucleotides represent locked residues.

### Bold nucleotides represent L-DNA residues.

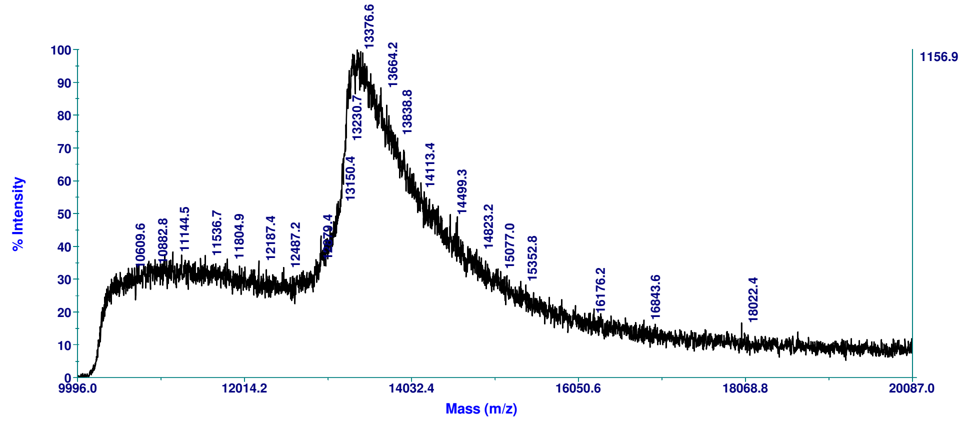


**Figure S1.** MALDI-TOF Mass Spectrometry analysis of EpCAM-L-DNA. [M]$^-$: 13376.6 (calc. 13267).

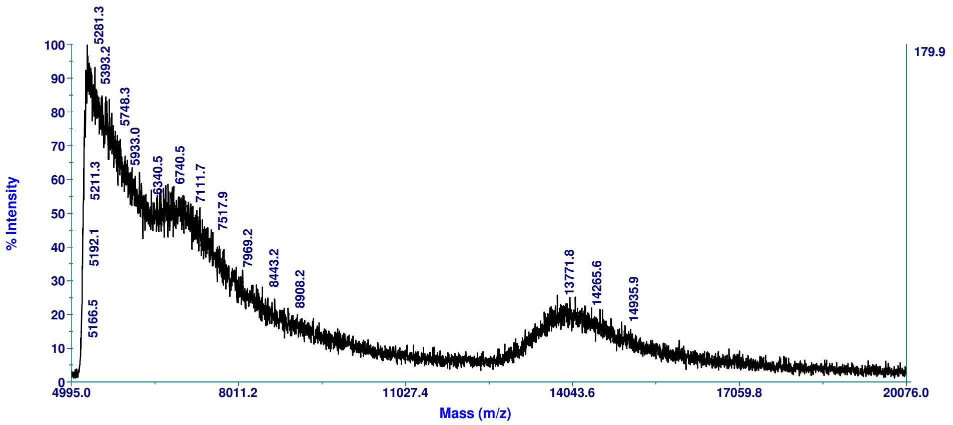


**Figure S2.** MALDI-TOF Mass Spectrometry analysis of cy5 labeled EpCAM-L-DNA. $[M]^-$: 13800 (calc. 13771.8).
